# Silicification of Cyanobacteria

**DOI:** 10.1101/2024.05.10.593471

**Authors:** Jun Sun, Yabo Han, Satheeswaran Thanagaraj

**Affiliations:** Research Centre for Indian Ocean Ecosystem, Tianjin University of Science and Technology, Tianjin 300457, China; Southern Marine Science and Engineering Guangdong Laboratory (Zhuhai), Zhuhai 519082, China; State Key Laboratory of Biogeology and Environmental Geology, China University of Geosciences, Wuhan, China; Institute for Advanced Marine Research, China University of Geosciences. Guangzhou, China

**Keywords:** Silicification, Cyanobacteria, Silaffins, Molecular clock, Algal evolution

## Abstract

The ability to use dissolved silicon has been found in various oceanic phytoplankton, including eukaryotic diatoms and prokaryotic cyanobacteria. While the silicification process in diatoms has been extensively studied, it is poorly known in cyanobacteria. In this study, the Cyanobacteria (*Synechococcus*) silicification and its evolutionary relationships of silicon-related proteins with different phytoplankton and algae were compared. Results showed that, compared to others, cyanobacteria have similar silicification proteins as diatoms. In detail, cyanobacteria and diatoms displayed three silicification-related proteins (SIT, Silaffins, and Pleuralins), while several genera showed one or two of them, suggesting that cyanobacteria might have developed a Si related mechanism earlier than the diatoms and other species. Our findings show that, despite cyanobacteria’s earlier silicon mechanism compared to others, they might not use efficient Si accumulation, perhaps adapting unique intracellular elemental variation for their cellular processes for growth potential.

## Introduction

Covering about 26% of the total elements present on Earth’s surface, silicon ranks second in terms of abundance **(Schneider and Boston, 1991)**. Majority of them can be located in the ocean as silicates with sources of continental weathering, biochemical sediments, and hydrothermal alteration **(Schneider and Boston, 1991)**. It was reported that dissolved silica (DSi) in the oceans was higher during the Precambrian period **(Conley and Carey, 2015; Conley et al., 2017)**, as determined by the equilibrium of mineral processes **(Schneider and Boston, 1991; Siever, 1992)**. For example, between the Ediacaran and Cambrian periods, the silicon concentration in the ocean was modified by siliceous sponges and radiolarians and changed the silicon deposition dynamics **(Conley et al., 2017; Maliva et al., 1989)**, that decrease in the concentration of DSi to approximately 1 mM **(Schneider and Boston, 1991; Tostevin et al., 2021)**. This concentration remained stable for ∼3 million years until the Mesozoic age and the Jurassic period, when diatoms emerged and proliferated **(Girard et al., 2020; Lee, 2018)**. As diatoms are very effective at precipitating DSi, the concentration of DSi in the oceans was quickly lowered to levels similar to the contemporary era **(Schneider and Boston, 1991)**. Therefore, diatoms play a pivotal role in the marine silicon cycle **(Armbrust, 2009)**, accounting for 20% of global carbon fixation **(Armbrust, 2009; Kroger and Poulsen, 2008)**.

Diatoms have been thought to dominate the marine silicon cycle **(Nelson et al., 1995)**, until a recent report suggested the biogenic silica (bSi) pool in the ocean may not entirely originate from diatoms and some associated with *Synechococcus* **(Wei and Sun, 2022)**. For example, *Synechococcus* makes up 20% of the particulate bSi in the euphotic zone in the Sargasso Sea, while in the equatorial Pacific Ocean, their contribution sometimes outweighed that of diatoms **(Baines et al., 2012)**. As *Synechococcus* are more abundant in open ocean, it was predicted that *Synechococcus* bSi stocks might be widespread **(Brzezinski et al., 2017). Tang et al. (2014)** have provided more evidence that *Synechococcus*-derived bSi standing stock accounts for 50% of the bSi inventory in the surface water, thus implying that half of the bSi in the surface water originates from *Synechococcus*. A recent study estimated that cyanobacteria responsible for 0.06–0.09 Tmol Si in the global ocean account for 2–3% of the diatom Si stock **(Wei et al., 2022)**.

Cyanobacteria are like diatoms in terms of DSi fixation **(Wei et al., 2022; Wei and Sun, 2022)**. A few measurements revealed that there are certain similarities between *Synechococcus* and the metabolism of Si in diatoms. Like diatoms, *Synechococcus* accumulates Si against a strong concentration gradient. According to **Baines et al. (2012)**, the total Si content of *Synechococcus* corresponds to internal values of 450 mM, while surface seawater concentrations are ∼10 μM in coastal waters and ∼1 μM in the open ocean. Such high intracellular concentrations would be comparable to those of diatoms of a size where amorphous silica would spontaneously precipitate from solution **(Iler, 1979)** and with soluble pools of Si between 19 and 340 mM **(Martin-Jézéquel et al., 2000)**. This is due to the fact that these *Synechococcus* are covered with an extracellular layer of polymeric substances (EPS), which encourage the precipitation of amorphous silica in the environment **(Handley et al., 2008; Tang et al., 2014; Moore et al., 2021)**. This EPS-Si produced by the *Synechococcus* is the precursor of the micro-blebs important to Si cycling **(Wei and Sun, 2022)**. In connection with this, **Tang et al. (2014)** suggested that intracellular Si can be deposited on extracellular polymeric substances in cyanobacteria, mediating silicon accumulation **(Moore et al., 2020)**, specifically in *Synechococcus* **(Baines et al., 2012; Brzezinski et al., 2017)**. Collectively, Si accumulation by cyanobacteria has provided a new dimension recently; however, their functional proteins or mechanisms for Si accumulation are still unclear.

It is known that diatoms use silicic acid transporters (SITs) **(Hildebrand et al., 1997; Durkin et al., 2016)**, Silaffins (proteins with silica affinity), and Pleuralins **(De Sanctis et al., 2016; Kröger and Wetherbee, 2000)** for their silicification. Briefly, SITs are the membrane protein that transports Si(OH)_4_ across the phosphate bilayer **(Thamatrakoln and Hildebrand, 2008)**, Silaffins are affinity proteins for silica **(Kroger and Poulsen, 2008)**, and Pleuralins are cell wall proteins coupled with silicon accumulation to cellular growth **(Kroger and Poulsen, 2008; De Sanctis et al., 2016; Kroger et al., 1997)**. Since cyanobacteria are like diatoms in terms of DSi assimilation, we hypothesize that *Synechococcus* may similarly silicify like diatoms-related Si proteins. Therefore, the purpose of this study is to detect and characterize these three proteins (SITs, Silaffins, and Pleuralins) in *Synechococcus* and other phytoplankton or algae and compare the evolutionary link between their occurrence and function using phylogenetic trees of molecular clocks.

## Materials and Methods

### Sequencing and phylogenetic tree construction

As diatoms are well known for their silicification, we used the UniProt database (Table S1) to investigate the existence of these SITs, Silaffins, and Pleuralins proteins in diatoms (*Cylindrotheca fusiformis* and *Thalassiosira pseudonana*). The obtained sequences were subsequently submitted to NCBI for evaluation. The number of algal species that included SITs, Silaffins, or Pleuralins was determined through our screening approach to be 42, 44, and 47 (Table S2), represents different phytoplankton or algae including Dinoflagellates, Cyanobacteria, Florideophyceae, Haptophyta, Xanthophyll, Charophyceae, Phaeophyceae, Chrysophyceae, Diatoms, Chlorophyceae, and Euglenophyceae. The 16S and/or 18S RNA sequences of each species were obtained from NCBI and used to construct phylogenetic trees using PhyloSuite **(Zhang et al., 2019)**.

The RNA sequences were aligned using MAFFT **(Katoh and Standley, 2013)**, and the tree-building model was identified using ModelFinder **(Kalyaanamoorthy et al., 2017)** (particularly, GTR+F+G4 was chosen for Silaffins and Pleuralins, whereas SYM+G4 was chosen for SITs). To accurately identify the fossil calibration sites during the molecular clock generation process, the phylogenetic trees were constructed using IQ-TREE **(Guindon et al., 2010; Minh et al., 2013; Nguyen et al., 2015)** and MrBayes **(Ronquist et al., 2012)** to reduce the margin of error in the trees.

### Time estimation of molecular clocks

Bayesian inference (BI) is a method of inferring statistics that uses the Bayes Rule of a particular hypothesis as more evidence and information. The maximum likelihood (ML) method is universal approach for estimating estimators. This ML method is representative of a class of phylogenetic tree reconstructions based on statistics and considers the probability of each nucleotide substitution in each sequence alignment. The Markov Chain Monte Carlo (MCMC) **(Metropolis and Ulam, 1949)** sequencing method was developed in the 1950s. It is a computer simulation method under the framework of Bayes Rule, where the sampling distribution changes with the simulation, with an effective calculation that guarantees the optimality of the solution theoretically.

In general, the evolution of different algal species was timed using the BEAST program **(Suchard et al., 2018)**. Briefly, the procedure required loading the (.nex) file into the Open BEAUti application and classifying each species of phytoplankton or algae according to the outgroup, ingroup, and cladistics of the phylogenetic tree in the Taxa module. The General Time Reversible (GTR) nucleic acid replacement model and the Gamma discrete rate distribution (with four rate classes) were then placed in the sites column of the BEAST evolution model. The relaxed molecular clock model was determined by the protocol described by **Drummond et al.’s (2006)** molecular clock model. In general, the evolution of different algal species was timed using the BEAST program **(Suchard et al., 2018)**. Briefly, the procedure required loading the (.nex) file into the Open BEAUti application and classifying each species of phytoplankton or algae according to the outgroup, ingroup, and cladistics of the phylogenetic tree in the Taxa module. The General Time Reversible (GTR) nucleic acid replacement model and the Gamma discrete rate distribution (with four rate classes) were then placed in the sites column of the BEAST evolution model. The relaxed molecular clock model was determined by the protocol described by **Drummond et al. (2006)** molecular clock model. The log.txt files were opened using the Tracer (http://tree.bio.ed.ac.uk/software/tracer/) software **(Rambaut and Drummond, 2007)** and verified the effective sample sizes (ESS) for each parameter. The results in general are considered credible if every ESS value is more than 200. However, if the execute (.xml) file is observed more than once or some of the ESS values are less than 200, this process should repeat. We followed this process for SITs, Silaffins, and Pleuralins, running 16, 13, and 25 MCMC analyses, respectively, to ensure accuracy. After that, we integrate the log.txt and trees.txt files for each independent MCMC analysis by setting the burnin value of 10%. We verified the ESS values of the log.txt file using Tracer and confirmed that all the values are above 200 to ensure convergence and an effective molecular clock evolutionary tree. We annotated the trees.txt file using TreeAnnotator by setting the burnin value to 10% for the output. We used FigTree v1.4.(http://tree.bio.ed.ac.uk/software/figtree/)to display the tree by selecting the *Node ages* from the *Display* in the *Node Labels* drop-down list and *height_95%_HPD* from the *Display* in the *Node Bars* drop-down list to show the 95% confidence interval.

## Results and Discussion

In this study, the protein sequences for SITs, Silaffins, and Pleuralins from *Cylindrotheca fusiformis* and *Thalassiosira pseudonana* were obtained from the UniProt database (Table S1) and are given in detail (refer to Supplementary Materials 2). It showed that *Synechococcus* SITs has high similarity with other species SITs, followed by Silaffins and Pleuralins. With these, the SITs, Silaffins, and Pleuralins sequences in *Synechococcus* were obtained by comparing these sequences to those in the NCBI database (Table S2). To examine these sequence results, phylogenetic trees were constructed using BI and ML methods. Interestingly, both methods produced trees with topological features that of similar and based on the molecular clocks for SITs, Silaffins and Pleuralins (Figures 1, 2, and 3).

Further, using the screening procedure that excluded cyanobacteria (including *Synechococcus*), Dinoflagellates, Phaeophyceae, Diatoms, Chlorophyceae, Euglenophyta, and Florideophyceae, with *Halobacterium salinarum* as the outgroup, a molecular clock phylogenetic tree for SIT was created.

The resulting tree demonstrates how different phytoplankton and algae have evolved over time, with Cyanobacteria emerging first, followed by Phaeophyceae, Euglenophyta, Florideophyceae, Dinoflagellates, Chlorophyceae, Chrysophyceae, and Diatoms (Figure 1). In similar, the evolution of Silaffins in different species was also analyzed using a phylogenetic tree based on the molecular clock method (Figure 2). It showed that Cyanobacteria, Florideophyceae, Dinoflagellates, Xanthophyll, Phaeophyceae, Diatoms, and Chlorophyceae were grouped together, representing the earliest to most recent stages of evolution. Our results are consistent with earlier observations of evolution, including cyanobacteria, dinoflagellates, and diatoms **(Lee, 2018)**.

**Figure 1.**
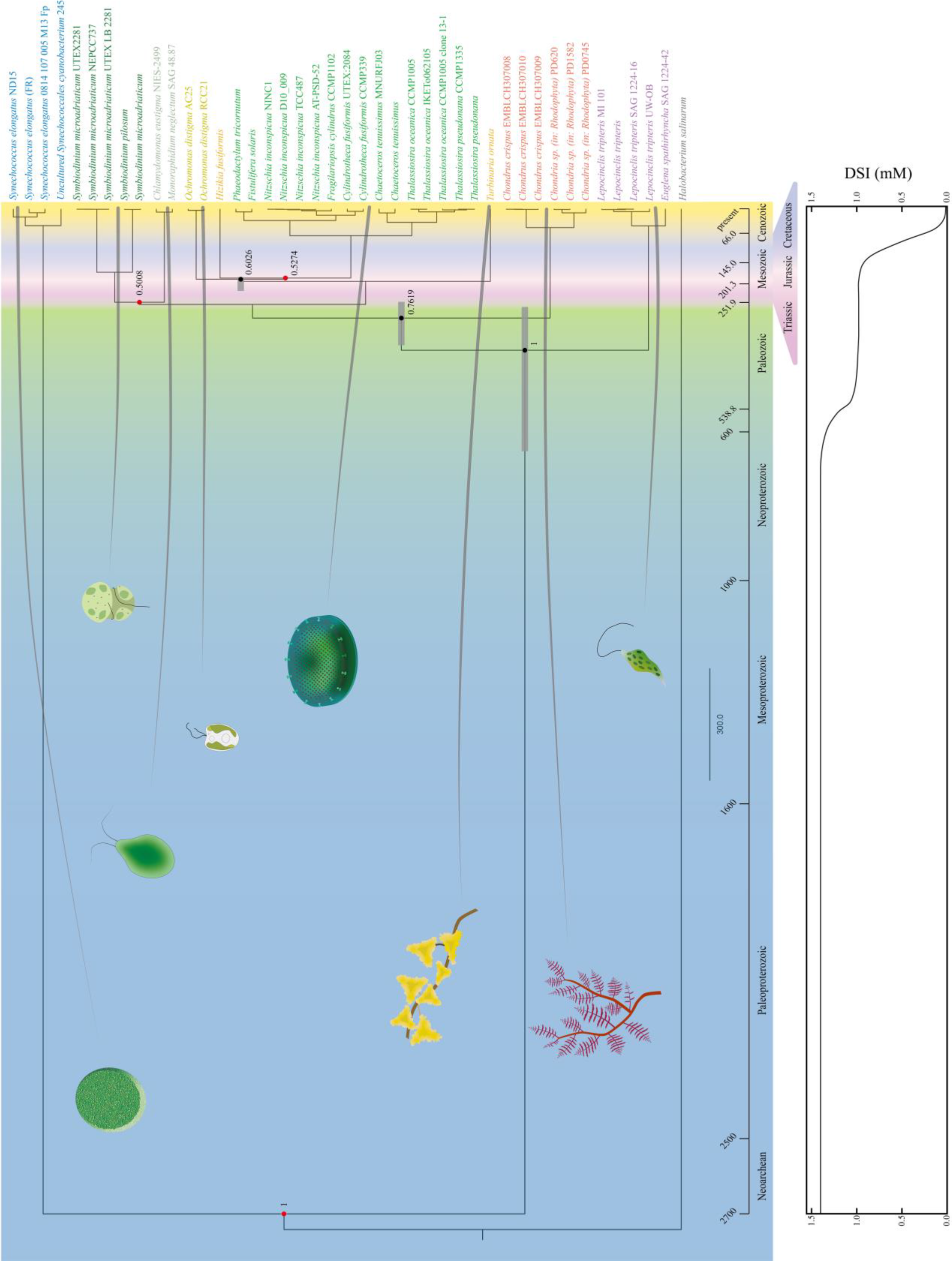
Divergence time and Marine silicon concentration for phytoplankton that may contain SITs-like or SITs -like proteins: Decimals near ancestral nodes indicate posterior probabilities. Solid red dots represent fossil calibration points, gray bar represents 95% confidence intervals, and solid black dots represent estimated divergence time by molecular clocks. 95% confidence intervals are shown here for only some nodes, Some solid points are not digitally labeled when their posterior probability is less than 0.5. Different colors in the background represent different geological periods. The diagram shows the representation of each class group. The graph below shows DSI concentrations of different geological ages, modified from **Conley et al. (2017)** and **Siever (1991)**.

**Figure 2.**
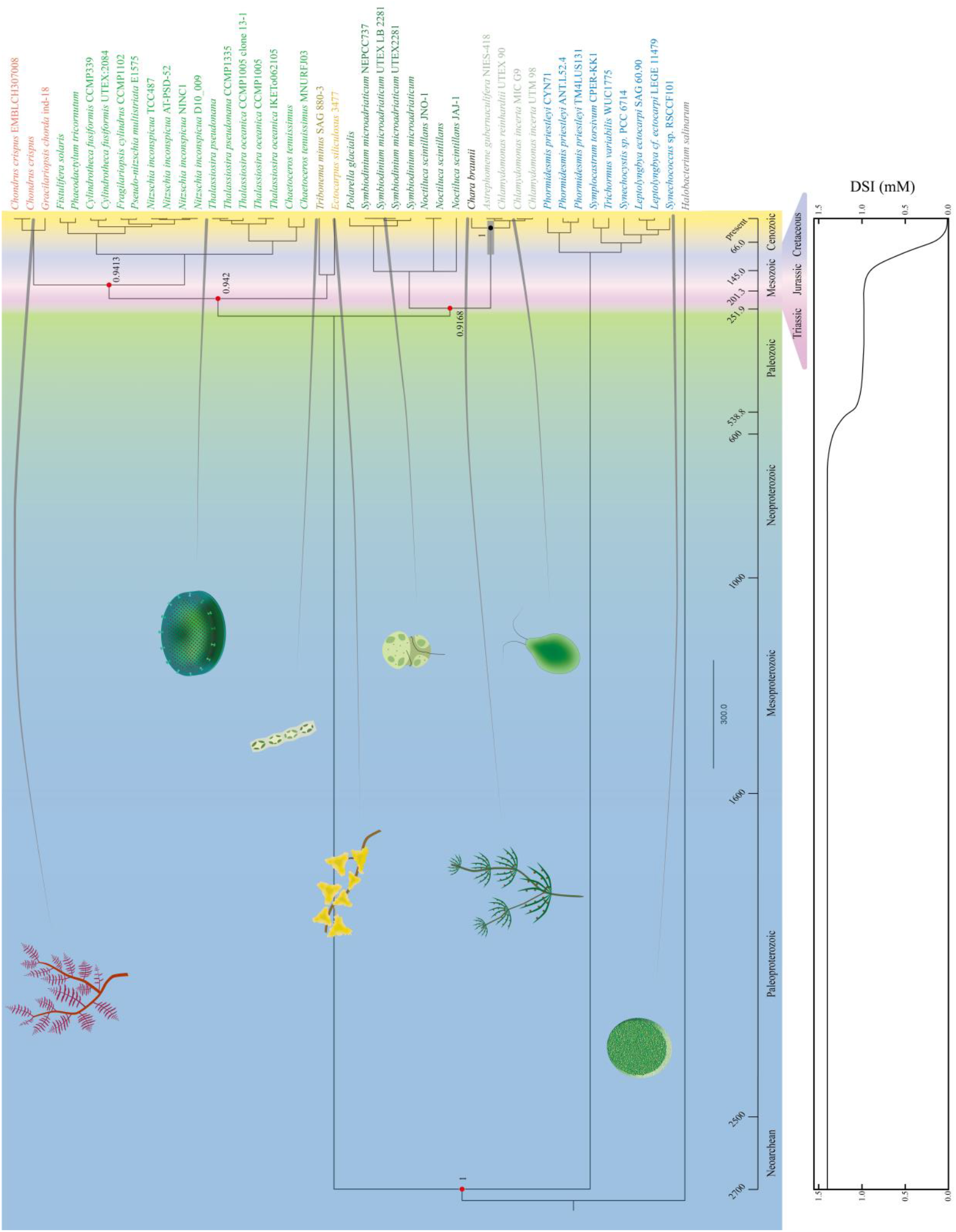
Divergence times and Marine silicon concentrations for phytoplankton that may contain Silaffins or Silaffins-like proteins: Decimals near ancestral nodes indicate posterior probabilities. Solid red dots represent fossil calibration points, gray bars represent 95% confidence intervals, and solid black dots represent estimated divergence times from molecular clocks, which represent 95% confidence intervals for only some nodes, Some solid points are not digitally labeled when their posterior probability is less than 0.5. Different colors in the background represent different geological periods. The diagram shows the representation of each class group. The graph below shows DSI concentrations of different geological ages, modified from **Conley et al. (2017)** and **Siever (1991)**.

Based on the earliest to the latest sequencing of Cyanobacteria, Dinoflagellates, Florideophyceae, Haptophyta, Phaeophyceae, Diatoms, and Chrysophyceae, the evolutionary relationship is demonstrated (Figure 3). It showed three sets of evolutionary changes (Supplementary Figure S1 & Table S3) represents 16 species only have SITs or their protein analogs; 13 species only have Silaffins or their protein analogs and 12 species only have Pleuralins or their protein analogs. In addition, one species showed both SITs and Silaffins or their protein analogs; 5 species have both SITs and Pleuralins or their protein analogs; 10 species have both Silaffins and Pleuralins or their protein analogs. Surprisingly, we noticed 20 species have all three proteins or their protein analogues (Supplementary Figure S1). The presence of these proteins and their analogues in cyanobacteria is given in brief (refer to Supplementary Table S3). In detail, one species of Cyanobacteria existed with SIT proteins or their analogues, eight species existed with Silaffin proteins or their analogues, and six species existed with Pleuralin proteions or their analogues. There were both SITs and Pleuralins, or their protein analogs, in three species of Cyanobacteria, and Silaffins and Pleuralins or their protein analogs, in one species of Cyanobacteria.

**Figure 3.**
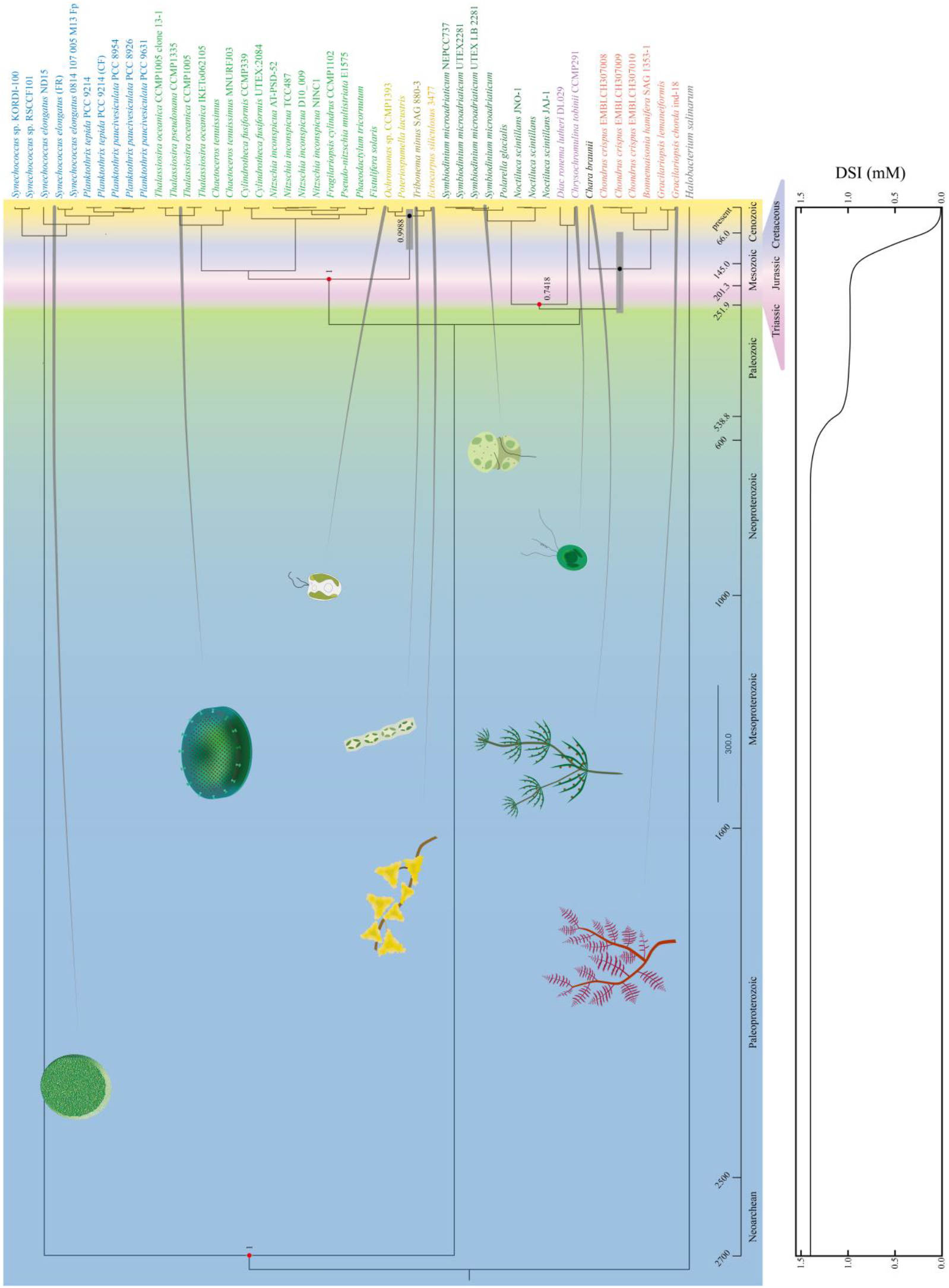
Divergence time and Marine silicon concentration for phytoplankton that may contain Pleuralins or Pleuralins-like proteins: Decimals near ancestral nodes indicate posterior probabilities.Solid red dots represent fossil calibration points, gray bar represents 95% confidence intervals, and solid black dots represent estimated divergence time by molecular clocks, which represent 95% confidence intervals for only some nodes, Some solid points are not digitally labeled when their posterior probability is less than 0.5. Different colors in the background represent different geological periods. The diagram shows the representation of each class group. The graph below shows DSI concentrations of different geological ages, modified from **Conley et al. (2017)** and **Siever (1991)**.

In the modern ocean, cyanobacteria make up a significant proportion of phytoplankton and are the only photoautotrophic prokaryotes currently in existence **(Falkowski et al., 2004)**. It is believed that clay can catalyze reactions **(Weiss and Hofmann, 1951)**, and silicon can be reflected in that silicon dioxide as a catalyst. Similarly, formaldehyde, ammonia, and hydrogen cyanide (HCN) can also be catalyzed to generate amino acids and even peptides on the surface of kaolin **(Wacker, 1958)**. Consistently, it is believed that silicate-related functions in living organisms today may be a development of the catalytic properties of silicon dioxide in the past **(Werner, 1977)**. Cyanobacteria are believed to have emerged during the Neoarchean era, approximately 2.7 billion years ago **(Lee, 2018)**. Six clones of *Synechococcus* experiment recently noticed a significant amount of silicon with growth rate unaffected by silicic acid concentrations between 1 and 120 μM suggesting *Synechococcus* lacks an obligate need for silicon. Their accumulation rates showed a bilinear response to increasing silicic acid from 1 to 500 μM with the rate of Si acquisition increasing abruptly between 80 and 100 μM **(Brzezinski et al., 2017)**. The notable alteration in the Si accumulation rate in *Synechococcus*, seen around 80–100 μM is comparable in the diatoms *Thalassiosira pseudonana* **(Thamatrakoln and Hildebrand, 2008)** and *Phaeodactylum tricornutum* **(Amo and Brzezinski, 1999)**.

It has been previously documented that silica accumulates in non-diatoms in a range of marine phytoplankton, such as cyanobacteria **(Bankston et al., 1979; Fisher et al., 1991)** and prasinophytes **(Fisher et al., 1991; D’Elia et al., 1979)**. While most of the phytoplankton evolution occurred during the Mesozoic era, cyanobacteria developed the ability to utilize dissolved silicon in the ocean during the Precambrian era. Throughout the Precambrian era, the density of phytoplankton evolution was relatively low, and the concentration of marine silicon remained at approximately 1.4 mM **(Conley et al., 2017)**, with amorphous silica saturation at 2 mM **(Iler, 1979; Krauskopf, 1956)**. As mentioned above, cyanobacteria lack an obligate need for Si with a concentration between 1 and 120 μM, and their notable alteration of Si accumulation has been recorded around 80-100 μM, similar to diatoms. These results lead us to speculate that during the Precambrian era, with a silicon concentration of 1.4 mM, cyanobacteria might have developed Si accumulation mechanisms earlier than that of diatoms; however, they did not efficiently used Si as like diatoms for their cellular processes; perhaps they balance the Si requirement for the intracellular elements for their growth potential. Our above assumption has been supported by prior research on the elemental variation of *Synechococcus* linked to physiological processes. According to **Van Mooy et al. (2009)**, investigation for example, if P is limited during *Synechococcus* growth, substituting S for P in phospholipids may result in a higher cellular S/P quota ratio. Subsequently, the elevated S/P ratio in *Synechococcus* cells may further improve cellular protein content and glutathione reductant concentrations (both enriched in N and S) for their growth **(Dupont et al., 2004; Twining et al., 2010)**. The key elemental behaviors, such as Si buildup or loss in *Synechococcus* cells, may be indirectly influenced by external variables through independent or strain-dependent physiological processes. This is because of the dynamic balance between cellular elements Si and C, P, and S **(Wei et al., 2022)**.

We screened the silicification genes and proteins in various phytoplankton and algae, including cyanobacteria, considering the apparent silicification of diatoms and the studies that have been done on them. Of them, cyanobacteria possess three silicification-related proteins (SITs, Silaffins, and Pleuralins), while others except dinoflagellate and Florideophyceae, have only one or two of these proteins. This further supports our assumption that Cyanobacteria tentatively inferred that the silicification function, which appeared early in Earth’s history, is relatively comprehensive and similar to that of diatoms and other species (Figure 4).

**Figure 4.**
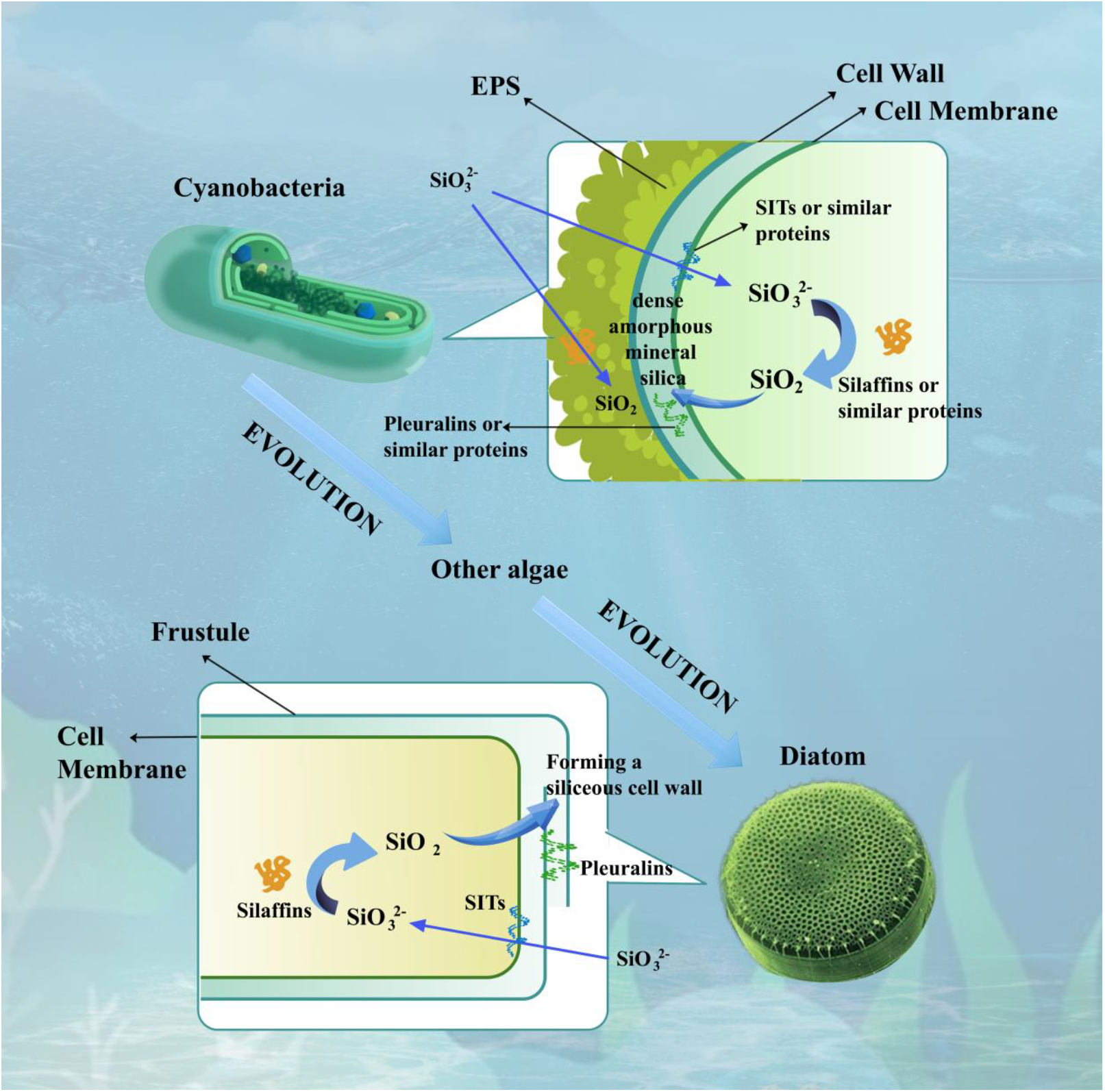
Since diatoms can accumulate silicon, it is speculated that Cyanobacteria can also accumulate silicon according to the evolutionary relationship. Cyanobacteria can transport silicates from the outside to the inside by SITs or similar proteins, and convert silicates into silica by Silaffins or similar proteins, The silica is then converted into dense amorphous silicon by Pleuralins or similar proteins that may exist in the periplasmic space between the cell wall and the cell membrane (Baines et al., 2012). At the same time, EPS may also contain Silaffins or similar proteins that can convert silicates in the Marine environment directly into silica (Tang et al., 2014).

Diatoms require large amounts of silicon to grow and form dense siliceous walls called frustules. Diatoms transport silicate from seawater around the cell into the cell by SITs, and Silaffins recognize and capture the silicates, convert them into silica, and transport them to the cell wall to combine with Pleuralins to form the cell wall. As a kind of Cyanobacteria, although the evolution time of *Synechococcus* is much earlier than that of diatom, it is still found that they have functional proteins related to silicate. **Ohnemus et al. (2016)** have measured the intracellular Si content of the North Atlantic and revealed that the intracellular Si content of *Synechococcus* varied greatly, ranging from 1 to 4,700 amol Si cell^−1^. **Brzezinski et al. (2017)** have reported that the growth rate of *Synechococcus* is not affected by the concentration of Si in culture. However, they also proposed that there are two Si pools, i.e., soluble and insoluble, in the cells of *Synechococcus* and speculated that soluble Si would likely bind to organic ligands. These results further support our assumption that *Synechococcus* might use their intracellular pools for their growth under Si variation.

Our results demonstrate the silicification of cyanobacteria, which is similar to the proteins related to silicon process of diatoms. Based on the results of previous studies on EPS, we speculated that EPS outside cyanobacteria cells may contain proteins similar to Silaffins, which can perhaps fix silicon in the marine environment so as to achieve the purpose of silicon accumulation. We also noticed that, similar to cyanobacteria, other phytoplankton or algae, i.e., Florideophyceae and Dinoflagellates, also conduct silicification processes, but the proteins associated with silication are not as diverse as the diatoms. It is due to the silicification process, which is sometimes absent or only partially present in the evolution of phytoplankton or algae. For instance, just a tiny fraction of the genes or proteins involved in the silicification process in the Charophyceae, Chrysophyceae, Xanthophyll, Phaeophyceae, and Chlorophyceae. This includes one or even two of the three types of genes involved in SITs (silicon input), Silaffins (silicon synthesis), and Pleuralins (silicon deposition). We assume that the intricate process of silicification arises from Cyanobacteria and involves the gradual loss of certain genes throughout the evolution of phytoplankton, ultimately resulting in their complete inheritance and enhancement by diatoms. However, further investigation should confirm these in more detail to better understand the phytoplankton influences in the marine silicon cycle.

This study aimed to explore the potential existence of three types of silicification proteins in cyanobacteria and various phytoplankton and algae to investigate their evolutionary relationships related to silicification. We noted that among all cyanobacteria, dinoflagellates, Florideophyceae, and diatoms have the three silicification-related proteins, while other species have one or two of these. Our results speculate that cyanobacteria might have developed a Si accumulation mechanism earlier than the diatoms and other phytoplankton, and they balance Si requirements from their intracellular elements for their growth potential. We understand that there are limitations on how the silicification process in *Synechococcus* is comparable to that of diatoms and its SIT (silicon intake), Silaffins (silicon synthesis), and Pleuralins (silicon deposition). However, due to the limitations of research on cyanobacteria in silicon utilization, future research should confirm the silicification process in different phytoplankton or algae and compare the silicification efficiency. In addition, the mechanism of silicification in different phytoplankton also requires a better understanding of the SITs, Silaffins, and Pleuralins in the silicification process, to better understand the marine Si cycle.

## Supporting information

Supplemental Table S1&2&3 and Figue S1

## Data availability

The data underlying this article are available in UniProt Database at https://www.uniprot.org/, and can be accessed with entry identifier. And NCBI Database in https://www.ncbi.nlm.nih.gov/, and can be accessed with accession number.

## Author Contributions

Conceptualization: Jun Sun; Methodology: Yabo Han; Writing-Original Draft: Jun Sun and Yabo Han; Writing-Review & Editing: Satheeswaran Thanagaraj. All authors reviewed the whole work and approved the final version of the manuscript.

## Acknowledgements

This research was financially supported by the Innovation Group Project of Southern Marine Science and Engineering Guangdong Laboratory (Zhuhai) (No. SML2022005).

