## Supplemental Table S1&2&3 and Figue S1 for "Silicification of Cyanobacteria"

**Silicification of cyanobacteria and its successors**

Supplementary tables

Supplementary table S1 The entry identifier for SITs, Silaffins, and Pleuralins in the Uniprot database.

| Protein Name | Entry Identifier | Species |
| --- | --- | --- |
| SITs | O81200 | *Cylindrotheca fusiformis* |
|  | O81199 | *Cylindrotheca fusiformis* |
|  | Q39510 | *Cylindrotheca fusiformis* |
|  | O81202 | *Cylindrotheca fusiformis* |
|  | O81201 | *Cylindrotheca fusiformis* |
|  | O81203 | *Cylindrotheca fusiformis* |
|  | B8C2V9 | *Thalassiosira pseudonana* |
|  | A5J0A3 | *Thalassiosira pseudonana* |
|  | B8C4N0 | *Thalassiosira pseudonana* |
|  | A5J095 | *Thalassiosira pseudonana* |
|  | B5YNJ0 | *Thalassiosira pseudonana* |
|  | A5J094 | *Thalassiosira pseudonana* |
|  | A5J093 | *Thalassiosira pseudonana* |
|  | Q0QVM8 | *Thalassiosira pseudonana* |
|  | Q0QVM6 | *Thalassiosira pseudonana* |
|  | Q0QVM7 | *Thalassiosira pseudonana* |
| Silaffins | Q9SE35 | *Cylindrotheca fusiformis* |
|  | B8BRK6 | *Thalassiosira pseudonana* |
|  | Q5Y2C2 | *Thalassiosira pseudonana* |
|  | Q5Y2C1 | *Thalassiosira pseudonana* |
|  | Q5Y2C0 | *Thalassiosira pseudonana* |
| Pleuralins | O22015 | *Cylindrotheca fusiformis* |

Supplementary table S2 16s/18s rRNA sequence Accession Number of each species in the NCBI database.

| Species Name | Accession Number |
| --- | --- |
| Synechococcus elongatus ND15 | AB871649.1 |
| Synechococcus elongatus (FR) | D83715.1 |
| Synechococcus elongatus 0814 107 005 M13 Fp | KP662909.1 |
| Uncultured Synechococcales cyanobacterium 245 | MW274389.1 |
| Symplocastrum torsivum CPER-KK1 | EF654065.1 |
| Trichormus variabilis WUC1775 | ON527149.1 |
| Phormidesmis priestleyi CYN71 | JQ687665.1 |
| Phormidesmis priestleyi ANT.L52.4 | AY493578.1 |
| Phormidesmis priestleyi TM4LUS131 | EU852504.1 |
| Synechocystis sp. PCC 6714 | AB041937.1 |
| Leptolyngbya ectocarpi SAG 60.90 | KC469578.1 |
| Leptolyngbya cf. ectocarpi LEGE 11479 | KU951732.1 |
| Phormidesmis priestleyi ULC062 | MH118732.1 |
| Synechococcus sp. RSCCF101 | MK471370.1 |
| Planktothrix tepida PCC 9214 | GQ351566.1 |
| Planktothrix tepida PCC 9214 (CF) | KU574121.1 |
| Planktothrix paucivesiculata PCC 8954 | GQ351576.1 |
| Planktothrix paucivesiculata PCC 8926 | GQ351574.1 |
| Planktothrix paucivesiculata PCC 9631 | GQ351578.1 |
| Synechococcus sp. KORDI-100 | KC192550.1 |
| Symbiodinium microadriaticum UTEX2281 | EF492514.1 |
| Symbiodinium microadriaticum NEPCC737 | EF492496.1 |
| Symbiodinium microadriaticum UTEX LB 2281 | KU900226.1 |
| Symbiodinium pilosum | M88518.1 |
| Symbiodinium microadriaticum | M88521.1 |
| Polarella glacialis | AF099183.1 |
| Noctiluca scintillans JNO-1 | AB297470.1 |
| Noctiluca scintillans | AF022200.1 |
| Noctiluca scintillans JAJ-1 | AB297469.1 |
| Ochromonas distigma AC25 | EF165136.1 |
| Ochromonas distigma RCC21 | KT860964.1 |
| Ochromonas sp. CCMP1393 | EF165142.1 |
| Poteriospumella lacustris | AY651074.1 |
| Hizikia fusiformis | AB011428.1 |
| Turbinaria ornata | JF718410.1 |
| Ectocarpus siliculosus 3477 | AY307398.1 |
| Phaeodactylum tricornutum | AY485459.1 |
| Fistulifera solaris | AB769957.1 |
| Nitzschia inconspicua NINC1 | AJ867021.1 |
| Nitzschia inconspicua D10_009 | FR873260.1 |
| Nitzschia inconspicua TCC487 | KC736636.1 |
| Nitzschia inconspicua AT-PSD-52 | KP662701.1 |
| Fragilariopsis cylindrus CCMP1102 | AY485467.1 |
| Cylindrotheca fusiformis UTEX:2084 | JN232981.1 |
| Cylindrotheca fusiformis CCMP339 | AY485457.1 |
| Chaetoceros tenuissimus MNURFJ03 | KT781055.1 |
| Chaetoceros tenuissimus | AB847417.1 |
| Thalassiosira oceanica CCMP1005 | HM991696.1 |
| Thalassiosira oceanica IKETo062105 | JX437375.1 |
| Thalassiosira oceanica CCMP1005 clone 13-1 | AF374479.1 |
| Thalassiosira pseudonana CCMP1335 | AY485452.1 |
| Thalassiosira pseudonana | AF374481.1 |
| Pseudo-nitzschia multistriata E1575 | KJ608081.1 |
| Chlamydomonas eustigma NIES-2499 | AB701493.1 |
| Monoraphidium neglectum SAG 48.87 | AJ300526.1 |
| Astrephomene gubernaculifera NIES-418 | LC086347.1 |
| Chlamydomonas reinhardtii UTEX 90 | AB511834.1 |
| Chlamydomonas incerta MIC G9 | JF834705.1 |
| Chlamydomonas incerta UTM 98 | KR349061.1 |
| Lepocinclis tripteris MI 101 | AF286210.1 |
| Lepocinclis tripteris | AF445459.1 |
| Lepocinclis tripteris SAG 1224-16 | AJ532456.1 |
| Lepocinclis tripteris UW-OB | AY935696.1 |
| Euglena spathirhyncha SAG 1224-42 | AJ532454.1 |
| Diac ronema lutheri DL029 | LC633056.1 |
| Chrysochromulina tobinii CCMP291 | KJ540196.1 |
| Chondrus crispus EMBLCH307008 | DQ316993.1 |
| Chondrus crispus EMBLCH307010 | DQ316997.1 |
| Chondrus crispus EMBLCH307009 | DQ316996.1 |
| Chondria sp. (in: Rhodophyta) PD620 | MF093919.1 |
| Chondria sp. (in: Rhodophyta) PD1582 | MF093921.1 |
| Chondria sp. (in: Rhodophyta) PD0745 | MF093920.1 |
| Chondrus crispus | Z14140.1 |
| Gracilariopsis chorda ind-18 | KF733542.1 |
| Bonnemaisonia hamifera SAG 1353-1 | AJ880421.1 |
| Gracilariopsis lemaneiformis | AY617149.1 |
| Chara braunii | AF032728.1 |
| Tribonema minus SAG 880-3 | AM490824.1 |

Supplementary table S3 The presence or absence of SITs, Silaffins, Pleuralins, or their protein analogues in different algae.

| SITs | Uncultured Synechococcales cyanobacterium 245 | MW274389.1 | Cyanobacteria |
| --- | --- | --- | --- |
|  | Symbiodinium pilosum | M88518.1 | Dinoflagellates |
|  | Ochromonas distigma AC25 | EF165136.1 | Golden Algae |
|  | Ochromonas distigma RCC21 | KT860964.1 | Golden Algae |
|  | Hizikia fusiformis | AB011428.1 | Brown Algae |
|  | Turbinaria ornata | JF718410.1 | Brown Algae |
|  | Chlamydomonas eustigma NIES-2499 | AB701493.1 | Green Algae |
|  | Monoraphidium neglectum SAG 48.87 | AJ300526.1 | Green Algae |
|  | Lepocinclis tripteris MI 101 | AF286210.1 | Euglenophyta |
|  | Lepocinclis tripteris | AF445459.1 | Euglenophyta |
|  | Lepocinclis tripteris SAG 1224-16 | AJ532456.1 | Euglenophyta |
|  | Lepocinclis tripteris UW-OB | AY935696.1 | Euglenophyta |
|  | Euglena spathirhyncha SAG 1224-42 | AJ532454.1 | Euglenophyta |
|  | Chondria sp. (in: Rhodophyta) PD620 | MF093919.1 | Red Algae |
|  | Chondria sp. (in: Rhodophyta) PD1582 | MF093921.1 | Red Algae |
|  | Chondria sp. (in: Rhodophyta) PD0745 | MF093920.1 | Red Algae |
| Silaffins | Symplocastrum torsivum CPER-KK1 | EF654065.1 | Cyanobacteria |
|  | Trichormus variabilis WUC1775 | ON527149.1 | Cyanobacteria |
|  | Phormidesmis priestleyi CYN71 | JQ687665.1 | Cyanobacteria |
|  | Phormidesmis priestleyi ANT.L52.4 | AY493578.1 | Cyanobacteria |
|  | Phormidesmis priestleyi TM4LUS131 | EU852504.1 | Cyanobacteria |
|  | Synechocystis sp. PCC 6714 | AB041937.1 | Cyanobacteria |
|  | Leptolyngbya ectocarpi SAG 60.90 | KC469578.1 | Cyanobacteria |
|  | Leptolyngbya cf. ectocarpi LEGE 11479 | KU951732.1 | Cyanobacteria |
|  | Astrephomene gubernaculifera NIES-418 | LC086347.1 | Green Algae |
|  | Chlamydomonas reinhardtii UTEX 90 | AB511834.1 | Green Algae |
|  | Chlamydomonas incerta MIC G9 | JF834705.1 | Green Algae |
|  | Chlamydomonas incerta UTM 98 | KR349061.1 | Green Algae |
|  | Chondrus crispus | Z14140.1 | Red Algae |
| Pleuralins | Planktothrix tepida PCC 9214 | GQ351566.1 | Cyanobacteria |
|  | Planktothrix tepida PCC 9214 (CF) | KU574121.1 | Cyanobacteria |
|  | Planktothrix paucivesiculata PCC 8954 | GQ351576.1 | Cyanobacteria |
|  | Planktothrix paucivesiculata PCC 8926 | GQ351574.1 | Cyanobacteria |
|  | Planktothrix paucivesiculata PCC 9631 | GQ351578.1 | Cyanobacteria |
|  | Synechococcus sp. KORDI-100 | KC192550.1 | Cyanobacteria |
|  | Ochromonas sp. CCMP1393 | EF165142.1 | Golden Algae |
|  | Poteriospumella lacustris | AY651074.1 | Golden Algae |
|  | Diac ronema lutheri DL029 | LC633056.1 | Haptophyta |
|  | Chrysochromulina tobinii CCMP291 | KJ540196.1 | Haptophyta |
|  | Bonnemaisonia hamifera SAG 1353-1 | AJ880421.1 | Red Algae |
|  | Gracilariopsis lemaneiformis | AY617149.1 | Red Algae |
| SITs+Silaffins | Thalassiosira pseudonana | AF374481.1 | Diatoms |
| SITs+Pleuralins | Synechococcus elongatus ND15 | AB871649.1 | Cyanobacteria |
|  | Synechococcus elongatus (FR) | D83715.1 | Cyanobacteria |
|  | Synechococcus elongatus 0814 107 005 M13 Fp | KP662909.1 | Cyanobacteria |
|  | Chondrus crispus EMBLCH307010 | DQ316997.1 | Red Algae |
|  | Chondrus crispus EMBLCH307009 | DQ316996.1 | Red Algae |
| Silaffins+Pleuralins | Synechococcus sp. RSCCF101 | MK471370.1 | Cyanobacteria |
|  | Polarella glacialis | AF099183.1 | Dinoflagellates |
|  | Noctiluca scintillans JNO-1 | AB297470.1 | Dinoflagellates |
|  | Noctiluca scintillans | AF022200.1 | Dinoflagellates |
|  | Noctiluca scintillans JAJ-1 | AB297469.1 | Dinoflagellates |
|  | Ectocarpus siliculosus 3477 | AY307398.1 | Brown Algae |
|  | Pseudo-nitzschia multistriata E1575 | KJ608081.1 | Diatoms |
|  | Gracilariopsis chorda ind-18 | KF733542.1 | Red Algae |
|  | Chara braunii | AF032728.1 | Charophyceae |
|  | Tribonema minus SAG 880-3 | AM490824.1 | Xanthophyll |
| All three protein | Symbiodinium microadriaticum UTEX2281 | EF492514.1 | Dinoflagellates |
|  | Symbiodinium microadriaticum NEPCC737 | EF492496.1 | Dinoflagellates |
|  | Symbiodinium microadriaticum UTEX LB 2281 | KU900226.1 | Dinoflagellates |
|  | Symbiodinium microadriaticum | M88521.1 | Dinoflagellates |
|  | Phaeodactylum tricornutum | AY485459.1 | Diatoms |
|  | Fistulifera solaris | AB769957.1 | Diatoms |
|  | Nitzschia inconspicua NINC1 | AJ867021.1 | Diatoms |
|  | Nitzschia inconspicua D10_009 | FR873260.1 | Diatoms |
|  | Nitzschia inconspicua TCC487 | KC736636.1 | Diatoms |
|  | Nitzschia inconspicua AT-PSD-52 | KP662701.1 | Diatoms |
|  | Fragilariopsis cylindrus CCMP1102 | AY485467.1 | Diatoms |
|  | Cylindrotheca fusiformis UTEX:2084 | JN232981.1 | Diatoms |
|  | Cylindrotheca fusiformis CCMP339 | AY485457.1 | Diatoms |
|  | Chaetoceros tenuissimus MNURFJ03 | KT781055.1 | Diatoms |
|  | Chaetoceros tenuissimus | AB847417.1 | Diatoms |
|  | Thalassiosira oceanica CCMP1005 | HM991696.1 | Diatoms |
|  | Thalassiosira oceanica IKETo062105 | JX437375.1 | Diatoms |
|  | Thalassiosira oceanica CCMP1005 clone 13-1 | AF374479.1 | Diatoms |
|  | Thalassiosira pseudonana CCMP1335 | AY485452.1 | Diatoms |
|  | Chondrus crispus EMBLCH307008 | DQ316993.1 | Red Algae |

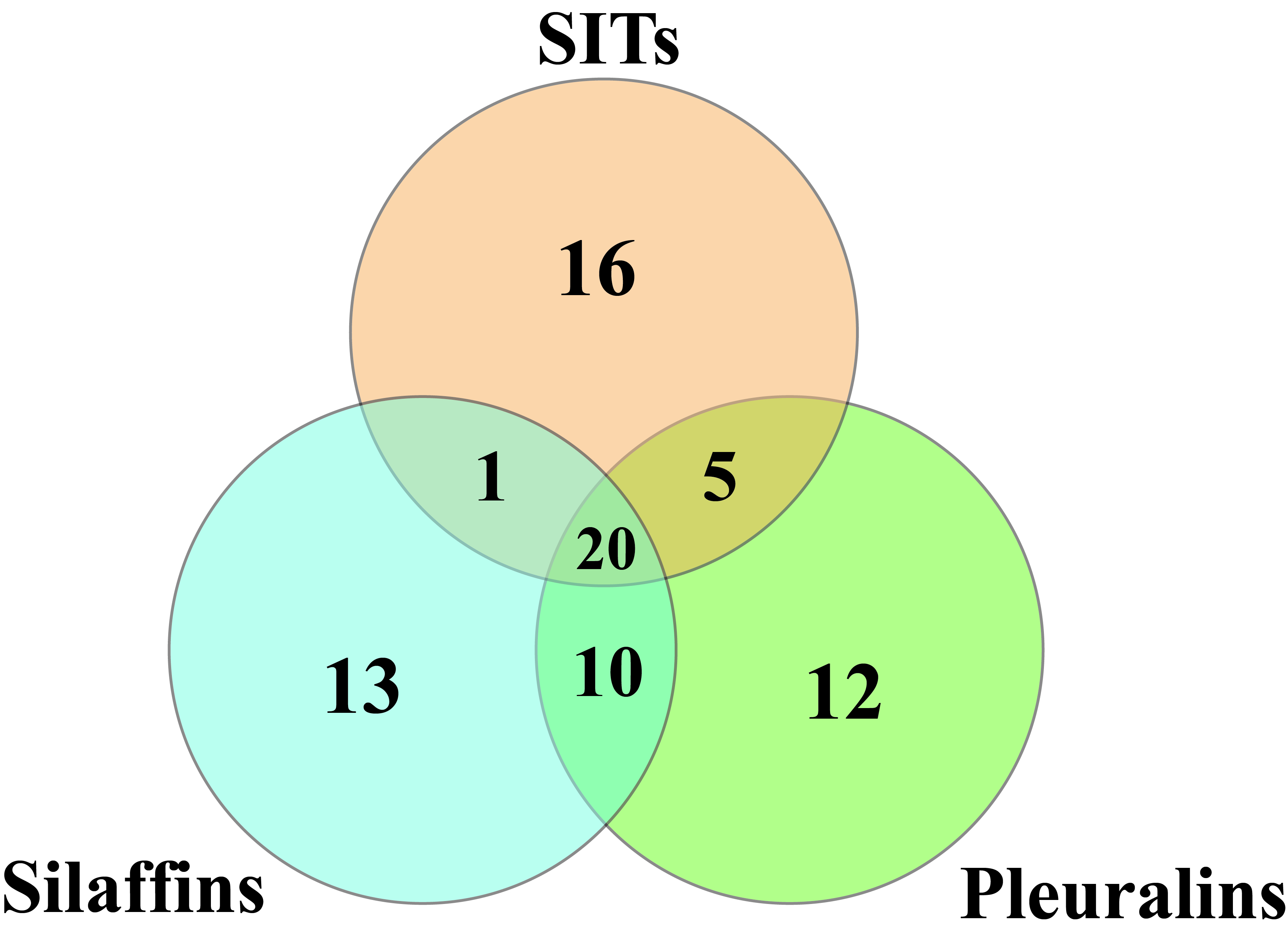

Supplementary figure S1 The distribution of SITs, Silaffins, Pleuralins, or their protein analogues in different algae. The number represents the amount of algae.
